# Functional basis of electron transport within photosynthetic complex I

**DOI:** 10.1101/2021.03.09.434417

**Authors:** Katherine H. Richardson, John J. Wright, Mantas Šimėnas, Jacqueline Thiemann, Ana M. Esteves, Gemma McGuire, William K. Myers, John J.L. Morton, Michael Hippler, Marc M. Nowaczyk, Guy T. Hanke, Maxie M. Roessler

## Abstract

Photosynthesis and respiration rely upon a proton gradient to produce ATP. In photosynthesis, the Respiratory Complex I homologue, Photosynthetic Complex I (PS-CI) is proposed to couple ferredoxin oxidation and plastoquinone reduction to proton pumping across thylakoid membranes, and is fundamental to bioenergetics in photosynthetic bacteria and some higher plant cell types. However, little is known about the PS-CI molecular mechanism and attempts to understand its function have previously been frustrated by its large size and high lability. Here, we overcome these challenges by pushing the limits in sample size and spectroscopic sensitivity, to determine arguably the most important property of any electron transport enzyme – the reduction potentials of its cofactors, in this case the iron-sulphur clusters of PS-CI, and unambiguously assign them to the structure using double electron-electron resonance (DEER). We have thus determined the bioenergetics of the electron transfer relay and provide insight into the mechanism of PS-CI, laying the foundations for understanding of how this important bioenergetic complex functions.

## Main Text

The majority of life on earth is dependent on photosynthesis, which uses light energy to generate potential energy in the form of a proton gradient. Transfer of electrons is coupled to movement of protons across a membrane by a set of exquisitely efficient molecular machines. Of these, the photosystems (PSII and PSI) and the cytochrome *b_6_f* complex have been well characterised, but until recently, little information was available about an additional proton pump, photosynthetic complex I (PS-CI, previously known as NDH-1). PS-CI is a key component of cyclic electron flow (CEF) in cyanobacteria and plants^1–4^. Photosynthetic organisms utilise CEF around photosystem I to increase the transmembrane proton gradient and thus ATP production, to meet the ATP:NADPH ratio required for CO_2_ fixation^5,6^. In addition to its role in cyanobacteria, PS-CI is also important in specific cell types that are common to some crop plants and critical for crop yield^7,8^. PS-CI accepts electrons from the terminal electron acceptor of PSI, ferredoxin (Fd), and reduces plastoquinone (PQ) (Figure 1a). This process is coupled to the pumping of protons and can therefore fine-tune the redox state of the compartment or cell under stress^9^.

**Figure 1:**
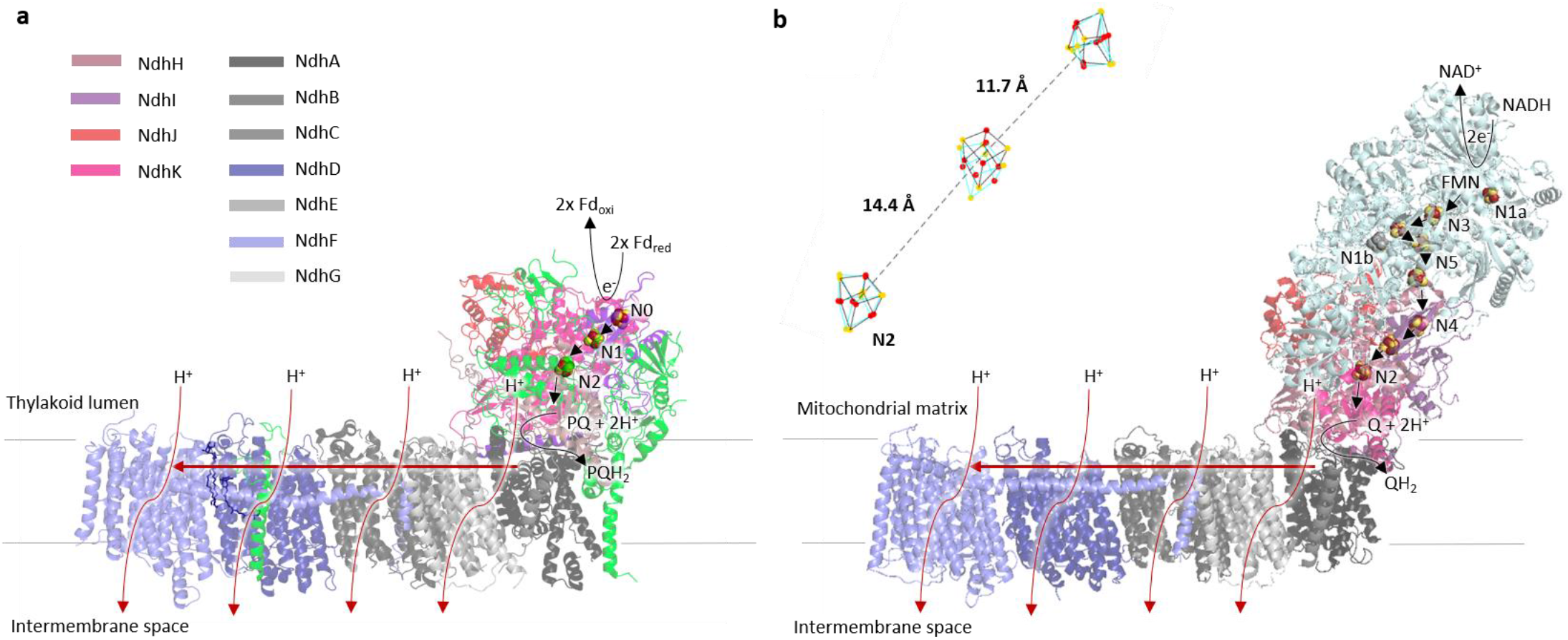
Structural and functional comparison of PS-CI and R-CI. **a** Structure and proposed catalysis of *T. elongatus* PS-CI (PDB: 6HUM). The 11 core subunits are coloured as indicated; the 7 oxygenic photosynthesis-specific (OPS) subunits are in green; reactions are shown schematically. Electron transfer from the donor Fd, to the acceptor PQ is indicated by black arrows. Putative movement of protons across (horizontal arrow) and through the membrane domain are indicated by red arrows. **b** Structure and proposed catalysis of *T. thermophilus* R-CI (PDB: 4HEA). The analogous subunits to PS-CI are coloured using the same key as in **a**, all other subunits are pale blue; reactions are shown as in **a**. FeS clusters in R-CI are labelled according to their EPR signals^15^. **b** Inset FeS clusters (Fe in red and S in yellow) of PS-CI (black, PDB: 6HUM) superimposed with structurally equivalent clusters in R-CI (cyan, PDB: 4HEA). Centre-to-centre PS-CI FeS cluster distances are labelled.

PS-CI was first identified as a homologue to respiratory complex I (R-CI)^10,11^. Recent cryo-EM structures have confirmed that PS-CI is a large multisubunit membrane protein with L-shaped architecture^12,13^. It comprises 11 core subunits and 7 oxygenic photosynthesis specific subunits (OPS) which are found in both hydrophobic and hydrophilic domains. The hydrophobic arm has four Mrp (Multiple-resistance- and-pH)-like Na^+^/H^+^ antiporters which likely translocate protons (Figure 1a)^2,14^. Although the core hydrophilic subunits are very similar in structure to their R-CI counterparts (Figure 1b), the hydrophilic domain is truncated by three subunits, including the NADH binding domain.

Electron transfer from NADH to ubiquinone through R-CI has been extensively studied; a non-covalently bound flavin mononucleotide transfers the 2 electrons from NADH singly down a series of 7 iron-sulphur (FeS) clusters to the ubiquinone binding site (Figure 1b)^16,17^. Although mechanisms have been suggested, how reduction of ubiquinone is coupled to proton pumping is not yet fully understood^18–20^. Electron paramagnetic resonance (EPR) spectroscopy has been a powerful technique to uncover critical information about the molecular environment of the FeS clusters and movement of electrons though R-CI. When reduced by its native substrate, NADH, up to five of the clusters can be observed in the EPR spectrum, with the other clusters remaining oxidised and therefore EPR silent^21^. The FeS cluster EPR signals have been named in order of their relaxation times (N1 > N2 > N3 > N4 > N5) and assigned to those identified in the structure^22,23^ (Figure 1b), revealing a ‘rollercoaster’ of alternating high and low reduction potential clusters, with the terminal [4Fe-4S] cluster N2 transferring electrons to ubiquinone^21,23–25^. Notably, although exact values vary and depend on pH, the N2 cluster is consistently more positive in reduction potential that the other FeS clusters^18,26,27^. In R-CI, cluster N2 is therefore postulated to act as an electron sink and may avert reverse electron transfer under physiological conditions, preventing backflow to O_2_ that would generate dangerous superoxide radicals via the reduced flavin^28^.

PS-CI accepts electrons from a one-electron donor, Fd, and is regulated by the OPS subunits unique to the photosynthetic complex^12,29^. Until recently, difficulties in purifying sufficient functional PS-CI have inhibited attempts to understand its function, and our knowledge has been limited to its subunit composition^9,30,31^, the phenotype of mutants and studies of its regulation using in vivo, or semi in vitro systems^32–34^. Thus, although several structures have recently been published due to advances in cryo-electron microscopy^12,13,29^, in contrast to R-CI, there is no experimental information about the molecular mechanism within PS-CI. In R-CI, EPR spectroscopic data, in particular information on FeS cluster reduction potentials^18,25^, preceded structural information by several decades, with pulse EPR later enabling a definitive assignment of the cluster properties to their spatial location in the electron transfer chain^35^. Although PS-CI was first discovered in 1998^36,37^, there is – perhaps surprisingly – no information on the reduction potentials of the electron transfer centres. Without reduction potentials it is extremely difficult to even formulate a hypothesis on how this molecular machine works. The lack of this most fundamental parameter for any electron transfer enzyme may not least be due to experimental bottlenecks, such as EPR-based potentiometric titrations and detailed pulse EPR measurements typically requiring very large amounts of enzyme, combined with the high magnetic anisotropy and extensive spin delocalisation of FeS clusters, whose EPR signals all overlap, making them one of the most challenging paramagnetic centres to work with. Here, we overcome these experimental bottlenecks and not only determine the first reduction potentials of the FeS clusters, but also assign their position in the electron transfer chain. We characterise the reduced FeS clusters of PS-CI from two strains of cyanobacteria using a combination of pulsed and continuous wave (CW) EPR spectroscopic methods. We determine the *g* values for all clusters and the reduction potentials of the two fully reducible clusters. Moreover, we provide conclusive assignment of thermodynamic properties to structurally defined FeS counterparts, giving novel insight into the functional mechanism of this crucial enzyme, placing it into the redox map of photosynthesis, and providing an essential foundation for future work on PS-CI.

To study the FeS clusters of PS-CI, the complex was purified from *Thermosynechococcus elongatus* using a native His-tag on NdhF1^13^, and from *Synechocystis* sp PCC6803 using a recombinant His-tag on NdhJ. The presence of the subunits was confirmed by proteomics (Supplementary Tables 1 & 2). The isolated complexes were reduced using sodium dithionite and suggest the presence of three reduced FeS cluster CW EPR signals (Figure 2a). Our recently reported high sensitivity EPR setup with a low-noise cryogenic preamplifier^38^ was employed to distinguish the overlapping FeS signals by performing different pulsed EPR relaxation filtering experiments^39^. Relaxation filtering selectively recovers the different FeS cluster spectra based on their spin-lattice and spin-spin relaxation times (Supplementary Figures 1 & 2 and Note 1). The N2 *g* values match very well with those for R-CI for both species of cyanobacteria^24,40,41^ (Supplementary Tables 3 & 4). Consistent with what is observed for R-CI, this FeS signal is observed at relatively high temperatures (20 K) and long relaxation times^39^. Given the structural and spectroscopic similarity between the species and R-CI N2, and in agreement with previous work on *T. elongatus* CI^12^, we assign the N2 EPR signal to the [4Fe-4S] cluster closest to the quinone binding site. Notably, an FeS cluster with these *g* values and with reduction potential −270 ± 25 mV was previously observed in EPR spectra of chemically reduced thylakoid membranes of two species of *Nostoc*, however its origin was unknown until now^42^.

**Figure 2:**
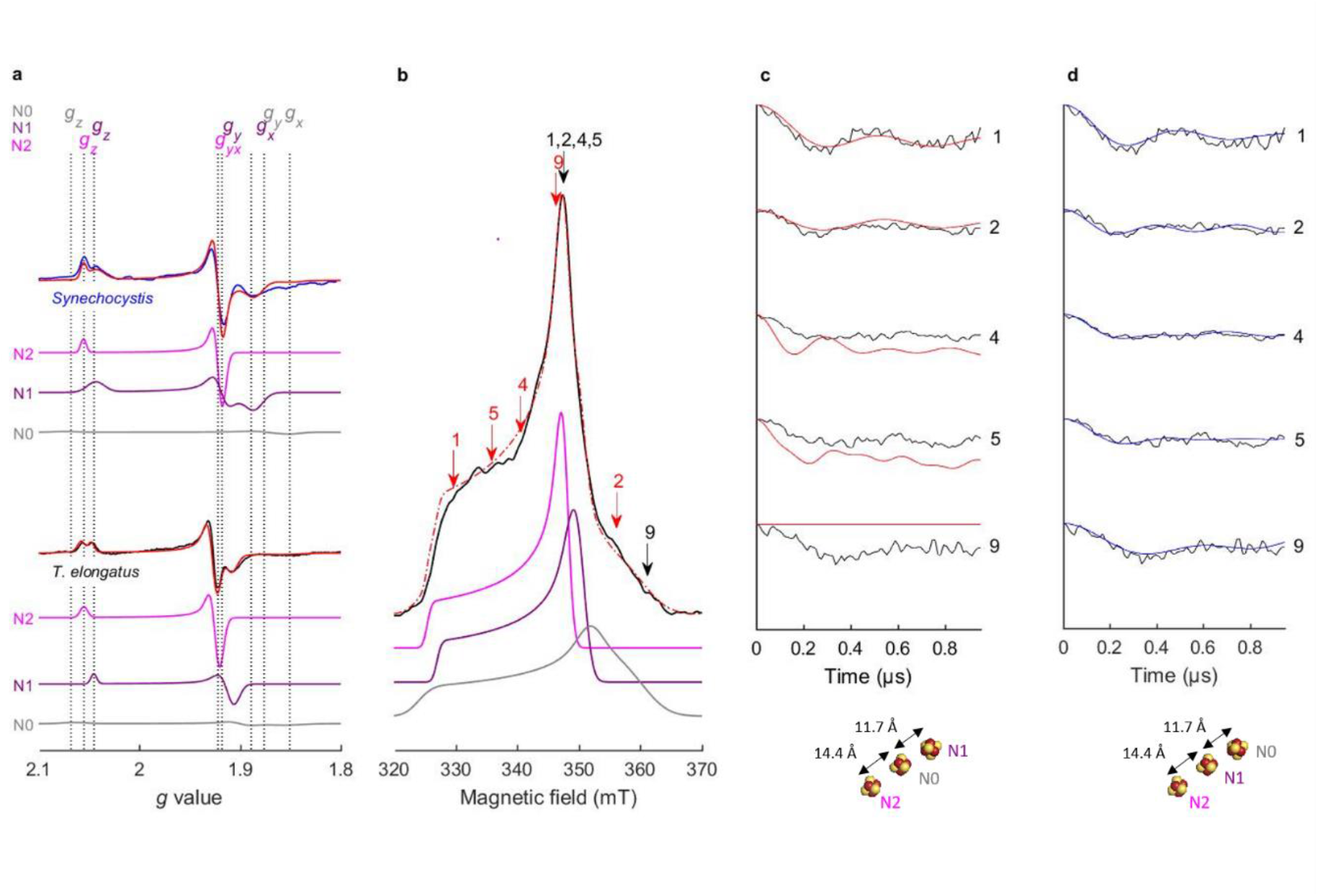
Assignment of the FeS cluster EPR signals to the structure of *T. elongatus* PS-CI using DEER. **a** CW EPR spectra (15 K) of FeS clusters in SDT reduced PS-CI of *Synechocystis* (blue) and *T. elongatus* (black). Simulations of the total (red) and individual N2 (pink), N1 (purple) and N0 (grey). *Synechocystis g* values: N2 (*g*_x,y_ = 1.922, *g*_z_ = 2.055), N1 (*g*_x_ = 1.886, *g*_y_ = 1.927, *g*_z_ = 2.043), N0 (*g*_x_ = 1.851, *g*_y_ = 1.867, *g*_z_ = 2.079) *T. elongatus g* values: N2 (*g*_x,y_ = 1.922, *g*_z_ = 2.055), N1 (*g*_x_ = 1.907, *g*_y_ = 1.913, *g*_z_ = 2.045), N0 (*g*_x_ = 1.852, *g*_y_ = 1.899, *g*_z_ = 2.064). See also Supplementary Table 3. **b** Set up of the pump pulse positions (red) and detection pulse position (black) for the corresponding DEER traces (for full experimental set up see Fig. S3). Echo-detected field sweep (black), sum of simulations (red), N2 (pink), N1 (purple), N0 (grey) (N2:N1:N0 ratio 1.00:0.92:0.90). **c,d** Orientation-selective DEER traces for the corresponding pump and probe positions (black). **c** Best-fit simulated DEER traces for model A, with the N1 cluster at 26.1 Å from N2 (red). **d** Best-fit simulated DEER traces for model B, with N0 at 26.1 Å from N2 (blue). Schematics of the structural models shown below. Note that the shorter distances (i.e. dipolar coupling to the middle cluster) do not contribute to the DEER traces (see Supplementary Note 2).

CW and pulsed EPR spectra of PS-CI in both species (Figure 2a, Supplementary Figures 1 and 2) could be well simulated (red traces) assuming two fully reduced [4Fe-4S] clusters and a third semi-reduced, which possess characteristic *g* values (Figure 2a). To decrease confusion with structural or spectroscopic [4Fe-4S] nomenclature for R-CI we refer to the remaining clusters as N1 and N0 for the reduced and semi-reduced clusters, respectively (Supplementary Note 1). The *g* values of the second fully reduced cluster N1 are similar between the two cyanobacterial species. However, in *Synechocystis* PS-CI N1 exhibits increased broadening, likely due to increased structural variation compared to *T. elongatus*^43^. N0 appears to be only partially reduced and its EPR signal is relatively broad. Although the *g* values of N1 and N0 are broadly consistent with those of other R-CI clusters, they do not match any one cluster well enough to assign them on the basis of homology (Supplementary Table 3 & 4). However, the PS-CI FeS cluster *g* values are similar between the photosynthetic species despite a wide evolutionary distance^44^; indicating that any heterogeneity between the complexes does not have a major effect on the FeS characteristics. Deconvolution of the three overlapping FeS signals in PS-CI through relaxation filtering provides the first unequivocal assignment of their *g* values.

To assign the respective PS-CI FeS cluster EPR signals to the clusters within the structure, we used double electron-electron resonance (DEER) spectroscopy (a pulsed EPR experiment that employs two microwave frequencies)^45^. The dipolar coupling between paramagnetic centres at the ‘pump’ and ‘probe’ microwave frequencies can be measured by analysing the modulation of the DEER spectra – this coupling strength is inversely proportional to the cubic distance between the centres providing structural information about the system (Supplementary Note 2)^46^. Multiple pump/probe positions that span the entire PS-CI EPR spectrum must be collected to calculate FeS cluster interaction distances (Figure 2b), as their highly anisotropic nature and limited bandwidth of microwave pulses results in a partial excitation of the EPR spectrum (orientation selection). The orientation selective DEER spectra were simulated with a custom program adapted from one previously developed for R-CI based on a local spin model^24^, taking into account the cluster positions (PDB:6HUM)^12^ and our experimentally determined *g* values (Figure 2c & d). With the position of N2 fixed, there are two possible models: model A, in which N2 and N1 are 26.1 Å apart, and model B in which N2 and the semi-reduced cluster N0 are 26.1 Å apart (Supplementary Figure 4). Only model B provides a good fit at all experimental pump and probe positions, both in terms of modulation frequency and depth. This is especially apparent at field position 9, where N1 does not contribute at the detection pulse position (Figure 2). We therefore assign the N0 signal to the [4Fe-4S] cluster adjacent to the Fd binding site.

Once the spectroscopic signatures of the clusters were assigned to structural positions, we determined the reduction potentials and therefore energetic favourability of electron transfer to and within PS-CI using small-volume potentiometric redox titrations (Figure 3)^47^. The EPR signal intensity of each cluster at each potential was estimated based on integration of the simulated spectra given the FeS cluster signal overlap (Figure 3, Supplementary Figure 5). The reduction potentials were estimated to be −220 and −230 mV ± 15 mV vs the standard hydrogen electrode (SHE) for N2 and N1, respectively, based on fitting the experimental data points to the one-electron Nernst equation. These values were consistent between cyanobacterial species. The reduction potentials are thus very similar and within experimental error not only between species, but also between the clusters (Figure 3). The clusters are therefore almost isopotential, meaning that electron transfer to N2 is as favourable as to N1.

**Figure 3:**
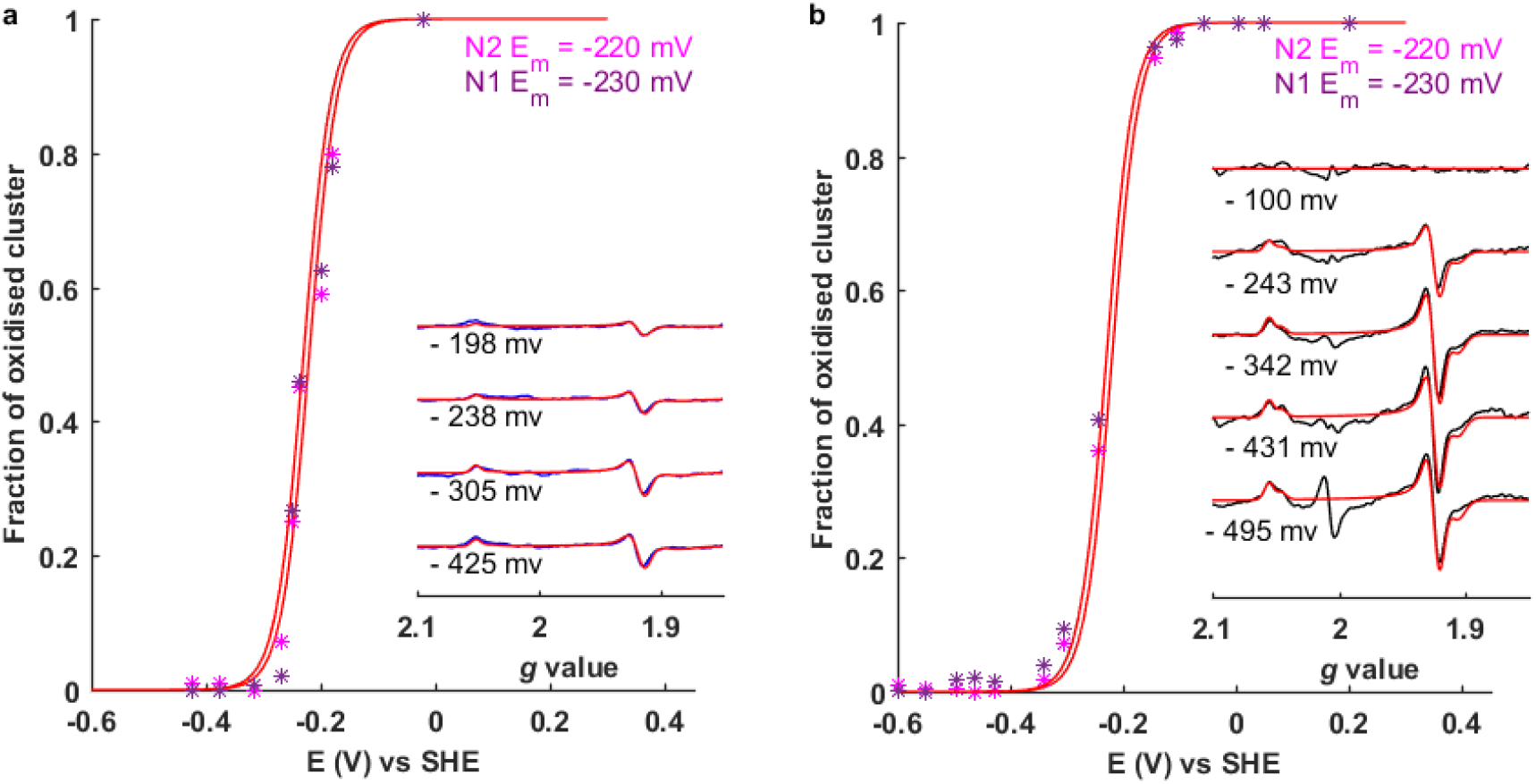
The reduction potentials of PS-CI N2 and N1. Small-volume potentiometric titrations of PS-CI from **a** *Synechocystis* and **b** *T. elongatus*. The fraction of oxidised cluster (N2 or N1) was determined from integration of the simulated CW EPR spectra (insets, simulations in red) normalised against fully reduced N2 or N1. Data points were fitted with the one-electron Nernst equation using the indicted midpoint potentials (*E*_m_).

The absence of a reduced N0 signal at −431 mV using measurement parameters that maximise N0 (Supplementary Figure 6) indicates that the reduction potential must be below ~-550 mV vs SHE. Such a low reduction potential of N0 will result in N2 and N1 being preferentially reduced, preventing backflow of electrons to form dangerous oxygen radicals in the cytosol. This is particularly important given the unknown contribution of PS-CI to free radical production, a process that initiates multiple defence and developmental signalling cascades in photosynthetic organisms^48,49^. R-CI is a notorious generator of the superoxide radical^50,51^, and it has recently been shown that blocking reverse electron transfer from the quinone site to the terminal flavin moiety prevents ROS production, protecting against cardiac ischemia-reperfusion injury^52^. PS-CI lacks this flavin cofactor, but the terminal FeS cluster is so solvent exposed that reverse electron transfer could also result in considerable free radical production^12,13,29^.

We were intrigued to find two clusters with equal and relatively positive *E*_m_ adjacent to each other as electrostatic repulsion would suggest this to be energetically unfavourable^53^. Moreover, our results are contrary to what is observed in R-CI and questions the mechanistic principle of alternating high- and low-potential clusters in electron transfer relays^21,54–56^. The redox potentials are very similar between the two species indicating this property is conserved. The isopotential nature of the clusters suggests that reduction of cluster N2 is unlikely to be involved in the reaction that couples electron transfer and proton translocation, in line with what is known about R-CI^57,58^. On the other hand, and contrary to R-CI, N2 and N1 in PS-CI may both be electron sinks, thereby facilitating the likely rate-limiting two-electron PQ reduction required for activity, from the one electron donor Fd.

In the cell, the very negative reduction potential of N0 would limit reduction of PS-CI until the Fd pool is in a highly reduced state (at least 100 times more abundant than PS-CI based on copy per cell estimates)^59,60^, with the close proximity of Fd to N0 providing an electron tunnelling pathway from Fd to N1/N2^29,53^. It remains possible that the potential is altered by the binding of subunits as is observed in R-CI^61^, or the substrate itself, as seen in photosystem I^62,63^. It has been suggested that electron transfer may occur in reverse in PS-CI if the proton motive force is high or the PQ pool is predominantly reduced, such as under decreased CO_2_ or fluctuating light intensities^2^. The very negative potential of N0 means that should such conditions occur, electron transport from N0 to Fd would be highly favourable, improving the capacity of Fd to effectively compete with oxygen as an oxidant and preventing oxidative damage. This finding provides a new basis for understanding under which cellular conditions the PS-CI complex is active, and how it could be intrinsically regulated by the redox status of the cell, necessitating physiological studies in cyanobacteria and higher plants.

The mechanism of function for both R-CI and PS-CI remain a matter of proposals that are still under debate, and although that of R-CI is much better understood, it is nonetheless unclear how electron transport is coupled to proton translocation across the membrane. The redox potentials provided in this work establish the energy gaps between the Fd electron donor and N0, between N0 and N1/N2, and between N2 and free quinone. This provides a platform for future work, in which new hypotheses can be tested regarding how electron transfer within PS-CI might be coupled to proton pumping. We have shown that overall electron transport to the quinone pool (at +80 mV^64^) is energetically very favourable in PS-CI. However there is no consensus on whether, or how, a quinone radical is stabilised in PS-CI^13,29,65^. Without accurate estimates of the reduction potential of PQ within PS-CI it is not yet possible to determine whether the coupling energy is indeed provided by quinol release, the latest theory-derived model suggested for PS-CI^66^, or quinone reduction.

Here we provide the first insight into the basis of the electron transfer mechanism in PS-CI, placing it into the bioenergetic network of photosynthesis and CEF in cyanobacteria. By applying custom EPR instrumentation and simulating the dipolar interactions of orientation dependent multi-coordinated paramagnetic species, the individual FeS cluster EPR signals and their corresponding structural position have been assigned using dilute (< 15 μM) and low-volume (~ 10 μL) samples. The reduction potentials of the N1 and N2 clusters allow overall favourable oxidation of reduced Fd and reduction of PQ, releasing the energy required for generating the proton gradient used by ATP synthase. We reveal the isopotential nature of the non-solvent exposed clusters, providing an energetic trap when Fd is not bound and theoretically preventing non-specific reverse electron transfer. These findings produce a first basis how PS-CI works in energetic terms, with the reduction potentials providing a solid platform for future studies to solve its mechanism of function and clarify its role in photosynthetic electron transport systems.

## Methods

### Purification of photosynthetic complex I

PS-CI was purified from *Synechocystis* sp. pcc 6803 by introducing a His-tag to the NDH-J subunit, large scale growth (70 litres of culture) and purification as described previously^67^. Briefly, cells were resuspended in 20 mL/L 20 mM sodium phosphate pH 7.5, 5% glycerol, 5 mM MgCl_2_, 10 mM NaCl, 1 mM benzamidine, 1 mM aminocaproic acid and 100 μM protease inhibitor. Cells were disrupted by passing twice though a microfluidiser (30,000 psi). Lysate supernatant was centrifuged at 50,000 g for 60 min at 4°C. Membranes were solubilised in 5 mm MgSO_4_, 20 mM MES pH 6.5, 10 mM MgCl_2_, 10 mM CaCl_2_, 25% glycerol (henceforth BB) with a final concentration of 1% (w/v) n-Dodecyl β-D-maltoside. The suspension was centrifuged at 50,000 g for 30 min 4°C. The supernatant was passed down a 25 mL Ni-NTA column equilibrated in BB (+ 10 mM imidazole, 0.03% DDM). PS-CI was eluted in BB supplemented with 200 mM imidazole and desalted by PD-10 column prior to EPR sample preparation. PS-CI was purified from *Thermosynechococcus elongatus* as previously described using a native His-tag on the NDH-F subunit^13^, prior to EPR sample preparation size exclusion chromatography was performed in 20 mM HEPES pH 8.0; 0.5 M mannitol; 150 mM NaCl; 0.03% (w/v) DDM. Both samples were concentrated in 100 kDa MWCO spin concentrators and PS-CI subunit presence confirmed using tandem mass-spectrometry.

### EPR sample preparation

EPR samples were prepared in a MBraun UniLab-plus glove box. 100 μL ~10 μM *Synechocystis* PS-CI in a 4.0 mm O.D. quartz EPR tube (Wilmad), and 10 μL ~20 μM *T. elongatus* PS-CI in a 1.6 mm O.D. Suprasil quartz EPR tube (Gross), were reduced in 20 mM sodium dithionite (Sigma, in Tris pH 9.5) for fully reduced spectra. The fully reduced samples were used for further pulsed EPR investigations (Figure 2).

Potentiometric titrations (Figure 3) on ~10 μM protein were carried out as previously described^47^. PS-CI was reduced or oxidised with substoichiometric amounts of sodium dithionite or K3Fe(CN)6 (Sigma) under anaerobic conditions in an electrochemical glass cell equipped with a 4°C water bath. Once equilibrated under nitrogen while stirring, 30 μM of the redox mediators methylene blue, indigotrisulfonate, indigodisulfonate, anthraquinone-2-sulfonate, benzyl viologen and methyl viologen (Sigma Aldrich) were added. The potential was measured using a Ag/AgCl mini-reference electrode (DRI-REF-2, World Precision Instruments) and a platinum working electrode (Scientific Glassblowing Service, University of Southampton; Pt from GoodFellow) and connected to an EmSTAT3+ potentiostat (PalmSens). Samples (~10 μL) were transferred to 1.6 mm O.D. Suprasil quartz EPR tubes (Gross) at the indicated potentials and flash-frozen in ethanol cooled from outside the glovebox by a dry ice acetone bath, before being transferred to liquid nitrogen. All reduction potentials are given relative to the potential of the standard hydrogen electrode (SHE). The reference electrode potential was determined to be +201 mV vs. SHE using quinhydrone (Sigma Aldrich) as an external standard.

## EPR measurements

### CW EPR spectroscopy

EPR measurements were performed using an X-band Bruker Elexsys E580 Spectrometer (Bruker BioSpinGmbH, Germany) equipped with a closed-cycle cryostat (Cryogenic Ltd, UK). All *T. elongatus* and titration measurements were carried out in an X-band split-ring resonator (ER 4118X-MS2). The field was calibrated using a DPPH standard (Bruker). The *Synechocystis* fully reduced sample was measured using an ER 4118X-MD5 resonator. Baseline spectra from samples containing only buffer or oxidised PS-CI were used as background and subtracted from the CW spectra. Unless otherwise specified data was collected at 15 K, 2 mW microwave power, 100 kHz modulation frequency, 7 G modulation amplitude and 16 scans.

### Relaxation filtered pulsed EPR spectroscopy

*T. elongatus* and *Synechocystis* samples were measured in an X-band split-ring (ER 4118X-MS2) and a dielectric ring (ER 4118X-MD5) resonators, respectively, both mounted on a modified standard EPR probehead containing a low-noise cryogenic preamplifier, which significantly enhances the EPR sensitivity^38^. Two-pulse echo-detected field sweeps (EDFS) were acquired with the pulse sequence π/2–*τ*–π–*τ*–echo. The *T*_1_-relaxation filtered EDFS were obtained using the sequence π -*T*_f_-π/2–*r*–π–*r*–echo, where *T*_f_ denotes the filtering time. Unless otherwise stated, the fully reduced *T. elongatus* PS-CI sample was measured at 10 K with π= 32 ns, *τ* = 250 ns, shot repetition time (SRT) of 2.04 μs. *Synechocystis* PS-CI was measured at 10 K, π = 32 ns, *τ* = 250 ns, SRT = 8.16 μs.

### DEER spectroscopy

DEER measurements used a four pulse sequence, π_A_/2–*τ*_1_–π_A_–(*τ*_1_+*t*)–π_B_–(*τ*_2_–*t*)–π_A_–*τ*_2_–echo^45^, with detection pulses at frequency ωA, and a single pump pulse at frequency ωB, which moved within the refocused echo sequence by variation of time *t*, e.g. steps of 12 ns. Experimental parameters for each position recorded are detailed in Supplementary Table 5. Position frequencies within the EPR spectrum are annotated in Supplementary Figures S3. An 8-step phase cycle was employed to remove unwanted echoes. Positions 1 to 5 were measured in ER 4118X-MS2, with *τ*_1_ = 134 ns and *τ*_2_ = 1.28 μs. Positions 6 to 9 were measured in ER 4118X-MS2 resonator equipped on a probehead containing a low-noise cryogenic preamplifier for increased sensitivity^38^, with *τ*_1_ = 400 ns and *τ*_2_ = 1 μs. All experiments were conducted at 10 K. DEER spectra were normalised to the zero-time intensity and to the scan number.

## Simulation of EPR spectra and analysis

CW and EDFS data was analysed and simulated with EasySpin esfit using Monte Carlo simulation in Matlab^68^. ‘Nernst plots’ (Figure 3) were generated based on the integrated area of the simulated signals (due to EPR signal overlap), plotted against the reduction potential of the samples, normalised to the maximum intensity signal resulting from the fully reduced sample. The one-electron Nernst equation was fitted to the experimental data points using the Matlab curve-fitting tool. The mid-point reduction potential error was calculated based on the 95% confidence intervals for the regression over the linear section of the Nernst curve. For details of DEER trace simulations see Supplementary Note 2.

## Acknowledgements

K.H.R. thanks the London Interdisciplinary Doctoral Programme for a studentship. The EPSRC (EP/T031425/1 to M.M.R., EP/L011972/1 to the Centre for Advanced ESR, University of Oxford and and EP/P510270/1 to M.Š.), BBRSC (BB/R004838/1 to G.T.H), John Fell Fund (0007019 to W.K.M.), DFG priority program 2002 (NO 836/4-1 to M.M.N.) and Leverhulme Trust (RPG-2018-183 to M.M.R.) are gratefully acknowledged for funding.

## Authorship contributions

K.H.R performed all data analysis and research except those specified below. G.T.H, M.M.R, K.H.R and designed the research with assistance from J.J.W and G.M. G.T.H and M.M.R directed the research. K.H.R and J.T. purified PS-CI from *T. elongatus* cells grown by J.T. and M.N. A.M.E cloned the NdhJ-His *Synechocystis* sp pcc 6803 strain. W.K.M performed DEER experiments 1-5. M.H performed the mass-spectrometry on PS-CI purified from *Synechocystis* sp pcc 6803. K.H.R and M.S. measured DEER traces 6-9 and the relaxation filtered EPR using the HEMT probe designed by M.S. and J.J.L.M. K.H.R, M.M.R and G.T.H wrote the manuscript. All authors read and approved the final manuscript.

