## Supplementary Information for "Functional basis of electron transport within photosynthetic complex I"

#### Supplementary Note 1. Assignment of the [4Fe – 4S] clusters of PS-CI using relaxation filtered EPR

The local environment of a paramagnetic species, and hence the extent of spin-orbit coupling of excited states into the ground state, causes deviation from the *g* = 2.0023 EPR signal of a free unpaired electron. Protein [4Fe-4S] clusters typically display axial or rhombic EPR signals with *g* values around 2.01 and 1.94 (where *g* values less than 2.0023 are the result of antiferromagnetic coupling between Fe centres). The exact *g* values, conformational strain and symmetry vary for each cluster, reflecting its particular environment. Clusters in similar structural environments have similar *g* values (Supplementary Table 4). PS-CI contains three [4Fe-4S] clusters which all contribute to some extent to the fully reduced spectra (Figure 2). The N2 cluster is structurally almost identical to R-CI (Figure 1) and therefore the *g* values are expected to be extremely similar in PS-CI. This is indeed the case (see Supplementary Table 2 & 3). The CW EPR spectra contain two fully reduced clusters, N2 and a second, defined here as N1. N0 is the third semi-reduced cluster that exhibits a very broad EPR signal (Figure 2). The three FeS cluster EPR signals overlap, making accurate assignment of their *g* values from CW EPR alone difficult.

As well as *g* values and strain, paramagnetic species have a spin-lattice relaxation time ($T_1$) and spin-spin relaxations time ($T_2$) influenced by their environment. Pulsed EPR was used to vary the contribution of different clusters to the EPR spectrum based on their relaxation times, by altering parameters of the applied microwave pulse sequence. The differing relaxation times can be used comparatively to filter the contribution of centres with distinct relaxation properties. Such 'relaxation-filtered' spectra were used to deduce the *g* values and strain for the three overlapping clusters, i.e. finding the values that fit each [4Fe-4S] cluster consistently (Supplementary Figures 1 & 2).

The first pulsed technique used to disentangle the FeS EPR signals in PS-CI was $T_1$-relaxation filtered EDFS[6]. In this experiment, an initial $\pi$ pulse inverts the spin magnetisation followed by a Hahn echo detection sequence applied after the filtering time $T_f$. By varying $T_f$, spin species having substantially different $T_1$ relaxation time can be resolved. PS-CI recovers N2 and N0 at long $T_f$ in both species (Supplementary Figure 1a & 2a). This indicates both N2 and N0 are recovered at relatively long $T_1$. N1 is recovered at intermediate $T_f$ (~8000 ns) indicating its $T_1$ is short compared to the other clusters.

Different measurement temperatures were also used to disentangle the overlapping spin species, which have different relaxation times. If a species is slow relaxing, it will begin to saturate at higher temperatures than faster relaxing species. As we decrease the temperature, the relative intensity of N1 and N0 increases compared to N2 for PS-CI from both species (Supplementary Figure 1b & S2b). N2 and N0 are also visible at higher temperatures than N1. This indicates N2 > N0 > N1 in $T_1$ relaxation time (i.e. N1 is the fastest relaxing cluster).

The above trend in $T_1$ relaxation time is further corroborated by the spin magnetization recovery, which depends on the shot repetition time (SRT) of the pulse EPR experiment. Slow relaxing species are not observed at short SRT due to saturation. The EPR signal intensity of N2 in *T. elongatus* PS-CI is reduced relative to N1 and N0 with decreasing SRT, therefore N2 is the slowest relaxing (Supplementary Figure 1c). N2 also decreases with shorter SRT in *Synechocystis* PS-CI, however the effect is much less pronounced (Supplementary Figure 2c).

Any species which has lost coherence during the spin evolution time $\tau$ will not be detected in an EDFS. By varying $\tau$, EPR FeS cluster signals are filtered primarily based on their spin-spin relaxation time $T_2$.

Signal N2 remains at long $\tau$ and is therefore the slowest relaxing in both species, consistent with its slow relaxing $T_1$ (Supplementary Figure 1d & 2d). N0 is filtered at long $\tau$ indicating that its $T_2$ is relatively short. The observation of a relatively long $T_1$ and short $T_2$ for N0 signifies a strong local magnetic field interacting with the cluster. This may be due to its proximity (~11 Å) to a fully reduced cluster (N1) or be an effect of inorganic ions present in the buffer given that N0 is almost solvent-exposed (based on the DEER assignment). N1 appears to also have a fast relaxation time ($T_1$ and $T_2$) in both species (Supplementary Figure 1d & 2d), consistent with its assignment of being the middle cluster, sandwiched between two paramagnets.

The differences in signal contribution afforded by varying these $T_1$ and $T_2$-dependent parameters enables a robust model of the *g* values and strains to be built for each induvial cluster (see Supplementary Table 3). The parameters thus determined yield good fits of all pulsed and CW spectra presented in this work. In the case of PS-CI, N2 has the slowest relaxation times; N1 has relatively fast relaxation times; and N0 has an unusual combination of slow $T_1$ with a fast $T_2$. To reflect the assignment of the EPR signals to their structurally defined clusters and to avoid confusion between the R-CI [4Fe-4S] clusters (N2, N3, N4 and N5) or [2Fe-2S] clusters (N1a or N1b), we have assigned the nomenclature N2, N1 and N0 to PS-CI.

#### Supplementary Note 2. Simulation of DEER spectra and analysis

An in-house programme that enables the simulation of orientation selective DEER spectra with highly delocalised spin centres, adapted from a previously reported programme[2], was used to simulate the DEER spectra. Briefly (further details below), all possible orientation selective DEER spectra were calculated for each possible assignment model and the resulting simulated spectra were compared with experimental data. The simulations take into account the experimental conditions used, the FeS cluster properties (cyroEM structure to obtain distances, *g* values and strain) and a local spin model for the spin Hamiltonian of [4Fe – 4S] clusters with spin projection factors $k_{1,2}$ =+1.17 $k_{3,4}$ =-0.67. Best fits to the experimental data were selected based on least-square fit residuals.

##### Simulation of orientation selective DEER to structurally assign the FeS clusters of PS-CI

Full accounts of the simulation algorithm[7], local spin model theory[8,9] and application to FeS cluster containing proteins[2] have been presented previously. The spin Hamiltonian for a pair of clusters (A and B) is written in terms of the intracluster spin-spin coupling interactions $H_{int}$, the Zeeman interaction $H_z$, the exchange interaction $H_{ex}$, and the dipolar interaction $H_{dd}$:

$$H = H_{int} + H_z + H_{ex} + H_{dd}.$$

The intracluster Hamiltonian ($H_{int}$ ) describes the interaction between the Fe spins, which couple to give an effective spin for the cluster; in clusters discussed here S = 1/2 is the ground state. The local spin model describes the individual spins of each Fe using the Wigner–Eckart theorem, where the spin column vector operator (**S**) for the *i*-th Fe in the cluster in cluster A (or B) is expressed in terms of the spin projection factor $k_i$:

$$\mathbf{S}_{Ai} = k_i \mathbf{S}_A$$

The Zeeman interaction ($H_z$) is calculated from the measured **g** matrices for the clusters, resulting from the weighted sum of the **g**-matrix contributions of each Fe center:

$$\mathbf{g_A} = \sum_i k_i \mathbf{g A_i}.$$

The exchange interaction ($H_{ex}$) is set to zero in our analysis as it is expected to be very small when the distance between the paramagnetic centres is >10 Å.

The dipolar frequencies observed in the DEER experiment result from the dipolar Hamiltonian:

$$\mathrm{H_{dd}} = \frac{\mu_0 \beta^2}{4\pi} \sum_i \sum_j \mathbf{S}_{\mathrm{A}_i}^{\mathrm{T}} \frac{\mathbf{g}_{\mathrm{A}_i}^{\mathrm{T}} \mathbf{g}_{\mathrm{B_j}} - 3\left(\mathbf{g}_{\mathrm{A}_i}^{\mathrm{T}} \mathbf{n}_{ij}\right)\left(\mathbf{n}_{ij}^{\mathrm{T}} \mathbf{g}_{\mathrm{B}_j}\right)}{r_{ij}^3} \mathbf{S}_{\mathrm{B}_j},$$

where **S** is the electron spin column vector operator, $r_{ij}$ - the distance, $\boldsymbol{n}_{ij}$ - the unit column vector. The **g** matrices used in the DEER simulations were experimentally determined, equivalent to an average value over the Fe subsites in the cluster. The averaging error introduced by this can be considered to be incorporated into the spin projection factors.

The dipolar coupling frequency, $\omega_{\mathrm{dd}}$, is approximated to[2,10]:

$$\omega_{\mathrm{dd}}^{m} = \frac{\mu_0 \beta^2}{2h} \mathrm{g}_{\mathrm{A}}^{m} \mathrm{g}_{\mathrm{B}}^{m} \sum_i \sum_j k_{\mathrm{A}_i} k\mathrm{B}_j \frac{3\cos^2 \psi_{ij}^{m} - 1}{r_{ij}^3},$$

where $\mathrm{g}_{\mathrm{A}}^{m}$ and $\mathrm{g}_{\mathrm{B}}^{m}$ are the experimental *g* values observed at the orientation of the unit magnetic field vector $B_0^m(\theta, \phi)$. The subscript *m* refers to a particular $(\theta, \phi)$ value for the unit magnetic field vector in the molecular frame. The simulation program samples $B_0^m(\theta, \phi)$ over half a unit sphere to include all possible orientations (as the spin Hamiltonian has inversion symmetry in the general case).

At X-band, FeS EPR spectra are significantly broader than microwave pulse excitation bandwidth, therefore only a subset of orientations, with respect to $B_0^m(\theta,\phi)$ will be excited by a microwave pulse. The DEER time trace contribution for orientation $B_0^m(\theta,\phi)$, where $f_{\mathrm{A}}^m$ of the A-spin spectrum at this orientation is excited is given by:

$$y(t)_{\mathrm{sim}}^{\mathrm{DEER},\,m} = f_{\mathrm{A}}^m\{(1-c_{\mathrm{mod}}^m) + c_{\mathrm{mod}}^m \cos(\omega_{\mathrm{dd}}^m t)\}$$

If $f_{\mathrm{A}}^m$ = 0, the orientation will not contribute to the simulation. If it is not, the intensity of $m$-th orientation with dipolar frequency $\omega_{\mathrm{dd}}$ weighted by $c_{\mathrm{mod}}^m$ and the nonmodulated part by $1 - c_{\mathrm{mod}}^m$. The modulation parameter, $c_{\mathrm{mod}}^m$, depends on the fraction B-spin resonance at $B_0^m(\theta,\phi)$ orientation that is excited by the pump pulse. Both these quantities are calculated by the integral between the A-spin (B-spin) spectrum at the orientation $B_0^m(\theta,\phi)$ and the position of the detection (pump) pulse profile.

The DEER traces are simulated for the experimental pump and detection microwave pulse positions (and relevant parameters) using the spin projection factors for ferredoxin-type FeS clusters (here $k_{1,2}$ = +1.17 $k_{3,4}$ = −0.67) that were previously found to be the suitable for R-CI[2], experimentally determined $g$ values, strain, and cluster coordinates from the published cryo-EM structure (PDB 6HUM)[1]. Simulations were generated for all possible orientations and weighted to the signal contribution from each cluster at the detection microwave position. Simulations were based on one interacting cluster pair (at a centre-to-centre distance of 26 Å, see Figure 1), plus decay due to relaxation from the third cluster. No significant difference in the fits was observed when all three potential interactions were taken into account in the simulations as the middle cluster is <15 Å from the other two, therefore the modulation frequency is too high to be observed with the DEER pulse sequence employed. The best-fit models are selected based on the residuals between the raw and simulated traces. The top 10% of fits were visually inspected to ensure the best fit was not a local minimum in the residual fits. None of the top 10% of fits for model A produced a better fit than those presented in Supplementary Figure 4. The $g$ tensors of the clusters that are the main contributors to the observed DEER spectra (N0 and N2, Model B) were found to be very similar to the R-CI N2 and N4 tensors reported previously[2].

#### Comparison with respiratory complex I

The experimentally observed modulation frequencies in the DEER spectra are similar to those previously reported for R-CI[2]. The simulations used to model the respiratory vs photosynthetic complex differ in two respects.

The first is the structure of N0 compared to N4 (both at ca. 26 Å from N2), as the cluster coordinates and principal $g$ values are different (Table S3 & S4) (whereas those of the respective N2 clusters are very similar). The overall cluster to cluster distances (taking centre-to-centre distances) only vary ~0.2 Å and hence does not have a profound effect on the modulation frequency. As the FeS cluster EPR signals are highly anisotropic, and the signals overlap, the extent to which principal $g$ values affect modulation of the DEER spectra depends on the pump and probe positions chosen and can be very minimal. The good fit for both models A and B at DEER position 1 (Supplementary Figure 4) exemplifies the importance of multiple pump and probe position to assign orientation selective DEER spectra. Hence, although the $g$ values of N4 and N0 differ slightly, their similar relative orientation (and hence distance) gives rise to similar orientation selective DEER spectra (for which pump/probe positions are comparable).

The second difference in the models is the calculation of modulation depth. In both cases simulated spectra are weighted to the signal intensity from individual clusters at different probe positions (as determined from EDFS, Figure 2b). In R-CI the cluster adjacent to N2 is not reduced, and therefore does not contribute to the DEER spectra. N4, N3 and N1b (a [2Fe 2S] cluster) all could have potentially contributed, and different models were calculated to reflect this, with N4 at 26 Å from N2 providing the best fit to experimental data. PS-CI contains only three FeS clusters which are all reduced to some extent. Therefore, all models were calculated using weights for both the clusters involved in the contributing distances. The accuracy of the simulated modulation depth and decay compared to experimental data, using the model for a partially reduced cluster N0, model B, confirms that the assignment is correct.

### Supplementary Tables

**Supplementary Table 1.** Tandem mass spectrometry of *T. elongatus* PS-CI.

| Subunit | ID | kDa | Protein score | Coverage (%) |
|---|---|---|---|---|
| NdhA | Q8DL32 | 41.3 | 25.44 | 22.96 |
| NdhB | Q8DMR6 | 55.1 | 20.15 | 8.74 |
| NdhC | Q8DJ02 | 13.7 | | |
| NdhD | Q8DKY0 | 56 | 138.31 | 21.21 |
| NdhE | Q8DL29 | 11.2 | 18.32 | 43.56 |
| NdhF | Q8DKX9 | 71.9 | 0.00 | 6.25 |
| NdhG | Q8DL30 | 21.6 | 117.93 | 25.50 |
| NdhH | Q8DJD9 | 45.2 | 344.24 | 63.20 |
| NdhI | Q8DL31 | 22.4 | 72.55 | 47.96 |
| NdhJ | Q8DJ01 | 19.2 | 66.88 | 64.29 |
| NdhK | Q8DKZ4 | 25.7 | 145.81 | 35.02 |
| NdhL | Q8DKZ3 | 11.6 | | |
| NdhM | Q8DLN5 | 12.6 | 444.83 | 49.55 |
| NdhN | Q8DJU2 | 8.3 | 68.95 | 79.33 |
| NdhO | Q8DMU4 | 7.86 | 110.44 | 78.57 |
| NdhP | | 4.9 | 0.00 | 36.36 |
| NdhQ | | 4.7 | | |
| NdhS | Q8DL61 | 8.1 | 80.61 | 62.16 |
| NdhV | | 13.6 | 17.50 | 55.20 |

**Supplementary Table 2.** Tandem mass spectrometry of *Synechocystis* PS-CI.

| Subunit | ID | kDa | Protein score | Coverage (%) |
|---|---|---|---|---|
| NdhA | P26522 | 40.5 | 55.40 | 24.5 |
| NdhB | P72714 | 55.4 | 38.43 | 15 |
| NdhC | P19045 | 13.7 | | |
| NdhD | P32421 | 57.5 | 113.01 | 18.5 |
| NdhE | P26524 | 11.5 | 16.95 | 13.6 |
| NdhF | Q55429 | 74.4 | 21.98 | 9.8 |
| NdhG | P26523 | 21.5 | 41.19 | 12.1 |
| NdhH | P27724 | 45.5 | 252.8 | 72.1 |
| NdhI | P36525 | 22.2 | 116.47 | 65.8 |
| NdhJ | P19125 | 20.6 | 69.58 | 41.3 |
| NdhK | P19050 | 27.3 | 112.69 | 81.5 |
| NdhL | P27372 | 9.3 | 2.72 | 12.5 |
| NdhM | P74338 | 14.1 | 59.75 | 57.9 |
| NdhN | P74069 | 17.6 | 115.16 | 72.7 |
| NdhO | P74771 | 8.3 | 38.51 | 63.9 |
| NdhP | | | | |
| NdhQ | | | | |
| NdhS | P74795 | 6.6 | 23.63 | 65.5 |
| NdhV | | | | |

**Supplementary Table 3**. Summary of FeS cluster g values and g strain in PS-CI. See Supplementary note 2 for details of the assignment. [a] Values taken from[1]

| | | *T. elongatus* [a] | *T. elongatus* | | *Synechocystis* | |
|---|---|---|---|---|---|---|
| | | *g* value | *g* value | *g* strain | *g* value | *g* strain |
| N2 | $g_x$ | 1.922 | 1.922 | 0.0092 | 1.921 | 0.0102 |
| | $g_y$ | 1.922 | 1.922 | 0.0112 | 1.921 | 0.0102 |
| | $g_z$ | 2.055 | 2.055 | 0.0087 | 2.055 | 0.0064 |
| N1 | $g_x$ | 1.899 | 1.907 | 0.0143 | 1.884 | 0.0204 |
| | $g_y$ | 1.911 | 1.913 | 0.0169 | 1.918 | 0.0158 |
| | $g_z$ | 2.045 | 2.045 | 0.0087 | 2.047 | 0.0200 |
| N0 | $g_x$ | 1.854 | 1.852 | 0.0376 | 1.851 | 0.0231 |
| | $g_y$ | 1.914 | 1.899 | 0.0201 | 1.867 | 0.0261 |
| | $g_z$ | 2.041 | 2.064 | 0.0271 | 2.07 | 0.0272 |

**Supplementary Table 4.** FeS cluster *g* values in R-CI from different species.

| | | ***B. taurus***[2] | ***Y. lipolytica***[3] | ***E. coli***[4] | ***T. thermophilus***[5] |
|---|---|---|---|---|---|
| N2 | $g_x$ | 1.921 | 1.925 | 1.903 | 1.93 |
| | $g_y$ | 1.927 | 1.929 | 1.903 | 1.94 |
| | $g_z$ | 2.054 | 2.053 | 2.05 | 2.048 |
| N4 | $g_x$ | 1.886 | 1.894 | 1.894 | 1.801 |
| | $g_y$ | 1.928 | 1.939 | 1.94 | 1.949 |
| | $g_z$ | 2.107 | 2.103 | 2.088 | 2.063 |

**Supplementary Table 5.** Experimental DEER parameters for the positions in Supplementary Fig. 3 and DEER traces in Supplementary Fig. 4.

| Position | Magnetic field (mT) | Detection frequency $\omega_A$ (GHz) | Pump frequency $\omega_B$ (GHz) | Detection pulse $\pi_A$ (ns) | Pump pulse $\pi_B$ (ns) |
|---|---|---|---|---|---|
| 1 | 398.9 | 9.116 | 9.676 | 20 | 14 |
| 2 | 352.6 | 9.484 | 9.316 | 24 | 12 |
| 2* | 352.4 | 9.484 | 9.316 | 24 | 10 |
| 3 | 346.0 | 9.350 | 9.616 | 24 | 16 |
| 4 | 344.1 | 9.256 | 9.574 | 24 | 12 |
| 5 | 344.1 | 9.256 | 9.665 | 24 | 14 |
| 6 | 350.8 | 9.512 | 9.359 | 12 | 20 |
| 7 | 357.4 | 9.588 | 9.361 | 20 | 18 |
| 8 | 356.9 | 9.360 | 9.590 | 16 | 18 |
| 9 | 357.0 | 9.302 | 9.590 | 16 | 18 |

### Supplementary Figures

**Supplementary Fig 1**. Relaxation filtering of the [4Fe – 4S] clusters of T. elongatus PS-CI. **a.** $T_1$-filtered ($\pi$ -$T_f$-$\pi$/2–$\tau$–$\pi$–$\tau$–echo) spectra of *T. elongatus* PS-CI (10 K), $\tau$ = 0.25 µs, filtration times ($T_f$, µs) indicated on the left **b.** Temperature dependence, **c.** shot repetition time (SRT, µs) dependence, and **d.** $\tau$ dependence (µs) of the Hahn echo ($\pi$/2–$\tau$–$\pi$–$\tau$–echo) of the FeS clusters signals of *T. elongates* PS-CI. Raw data in black, sum of simulations in red. Percentage contributions of N2 in pink, N1 in purple and N0 in grey.  See supplementary note 2 for more details.

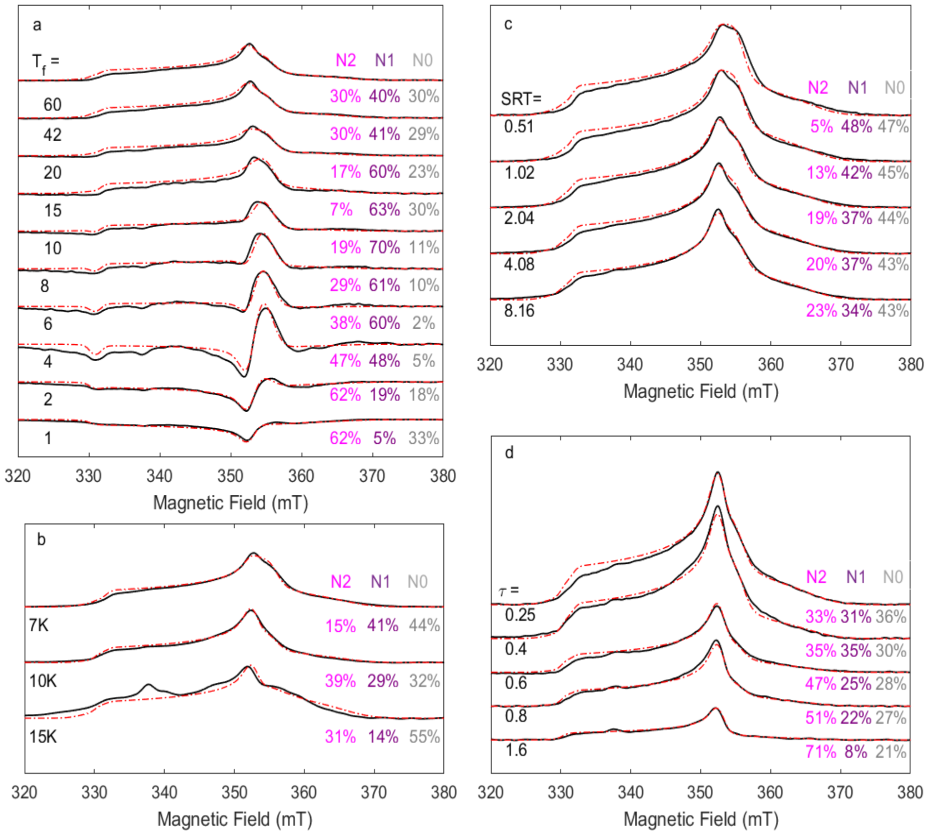

**Supplementary Fig 2.** Relaxation filtering of the [4Fe – 4S] clusters of Synechocystis PS-CI. **a.** $T_1$-filtered ($\pi$ -$T_f$- $\pi/2$–$\tau$– $\pi$ –$\tau$–echo) spectra of *Synechocystis* PS-CI at (10 K), $\tau$ =0. 25 µs, filtration times indicated on the left ($T_f$, µs) **b.** Temperature dependence, **c.** shot repetition time (SRT, µs) dependence, and **d.** $\tau$ dependence (µs) of the Hahn echo ($\pi/2$–$\tau$–$\pi$–$\tau$–echo) of the FeS clusters signals of *Synechocystis* PS-CI. Raw data in black, sum of simulations in red. Percentage contribution of N2 in pink, N1 in purple and N0 in grey. The peak observed at about 345 mT in the *Synechocystis* spectra is a $g$ = 2 signal originating from an organic radical. See supplementary note 2 for more details.

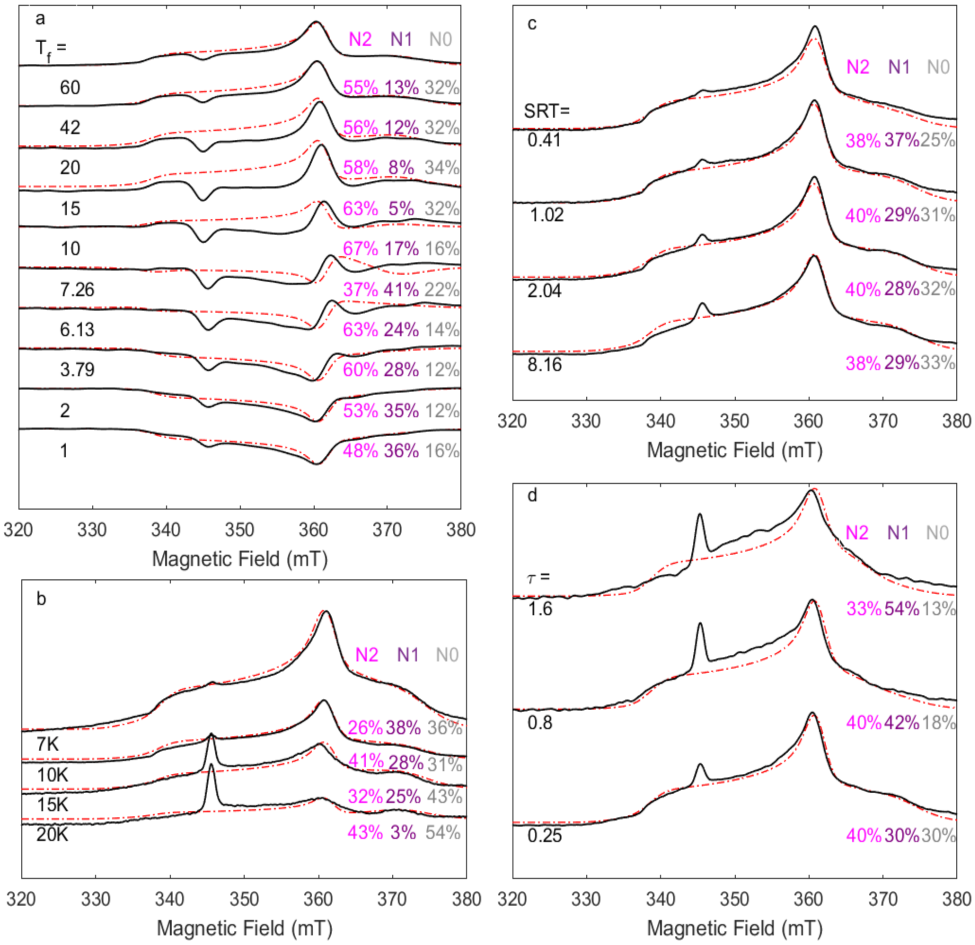

**Supplementary Fig 3.** DEER pump and detection pulse set up. **a.** Representation of the orientation selection of the microwave pulses for position 2 **b**. The pump pulse excitation profile for a 12 ns $\pi$-pulse at frequency $\omega_B$ = 9.316 GHz (red) and the detection pulse excitation profile for a 24 ns $\pi/2$-pulse at $\omega_A$ = 9.484 GHz (black) applied to *T. elongatus* PS-CI. Only a fraction of each spectrum is excited by the microwave pulses. The fixed field position of the DEER measurement is at 352.6 mT. **b.** Pump pulse positions in red, detection pulse position in black corresponding DEER traces in Supplementary Fig. 4. Full experimental parameters in Supplementary Table 5, raw data in black, sum of simulations in red, N2 in pink, N1 in purple, N0 in grey (N2:N1:N0 ratios are 1.00:0.92:0.90).

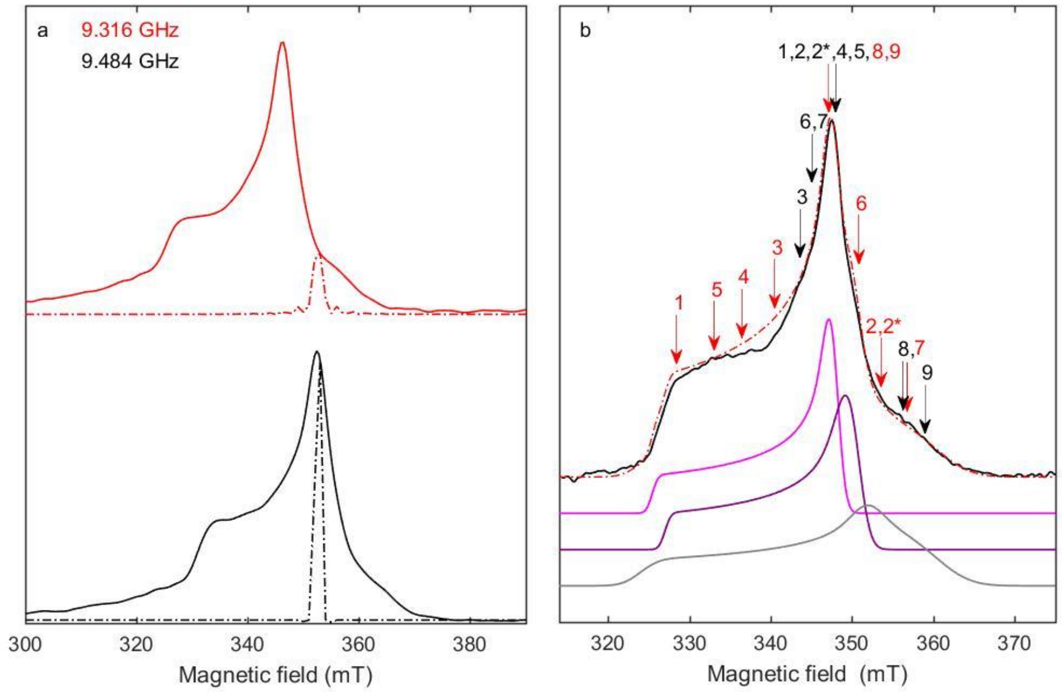

**Supplementary Fig 4.** Complete set of experimental DEER spectra of PS-CI and best-fit simulations for both models. DEER spectra from *T. elongatus* measured at 10 K, magnetic field (mT), detection ($\omega_A$, GHz) and pump pulse ($\omega_B$, GHz) positions (corresponding to the pump and detection pulse positions shown in Supplementary Fig. 3), detection $\pi$ length ($\pi_A$, ns) and pump $\pi$ length ($\pi_B$, ns) listed in Supplementary Table 5; raw data in black. The models correspond to the cluster order indicated below, distances calculated from the intercluster coordinates of PDB:6HUM. All traces are normalised to the intensity at zero time and the scan number.

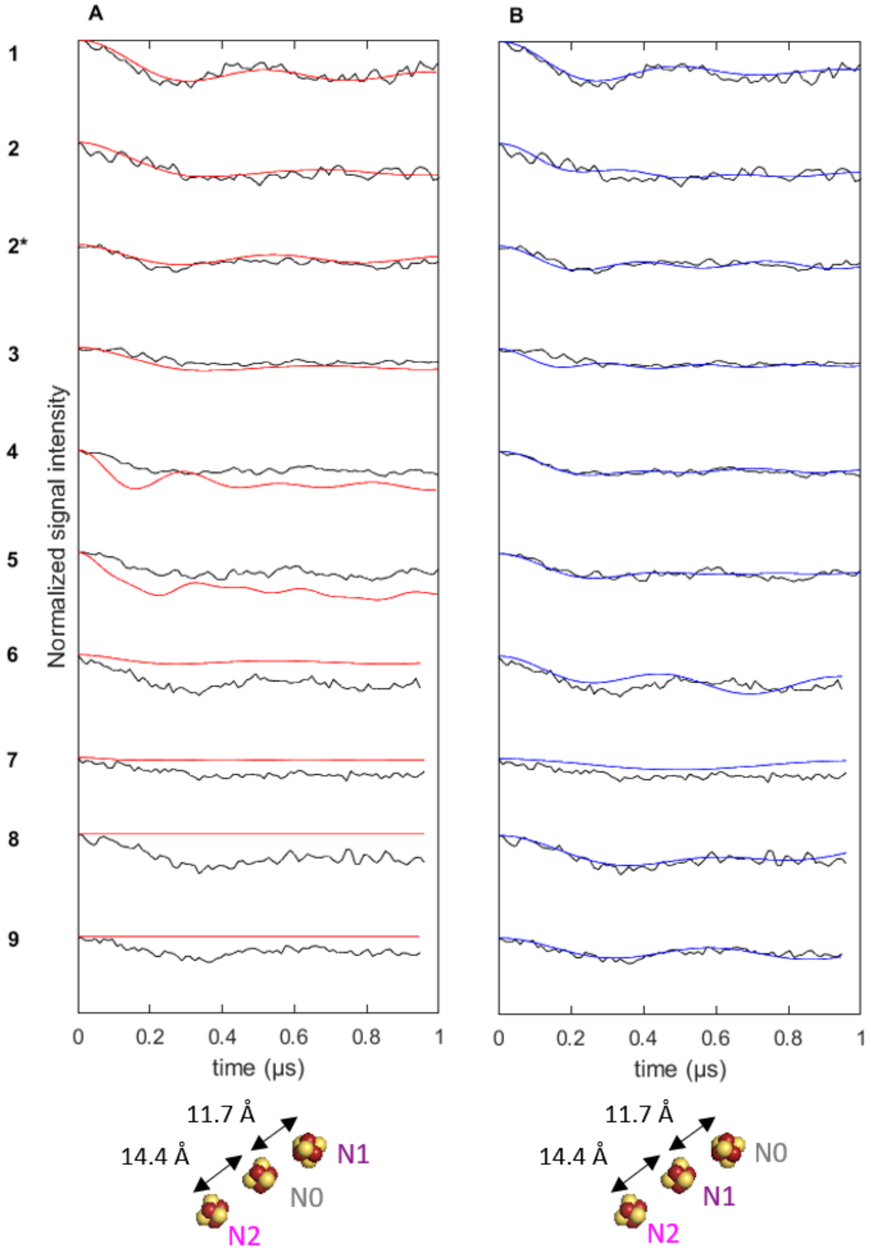

**Supplementary Fig 5**. EPR-based Potentiometric Redox Titrations of **a.** *T. elongatus* and **b.** *Synechocystis* PS-CI. The reduction potentials (mV) of the samples are indicated on the right-hand side. Raw data in black and simulations in red. SDT = sodium dithionite.

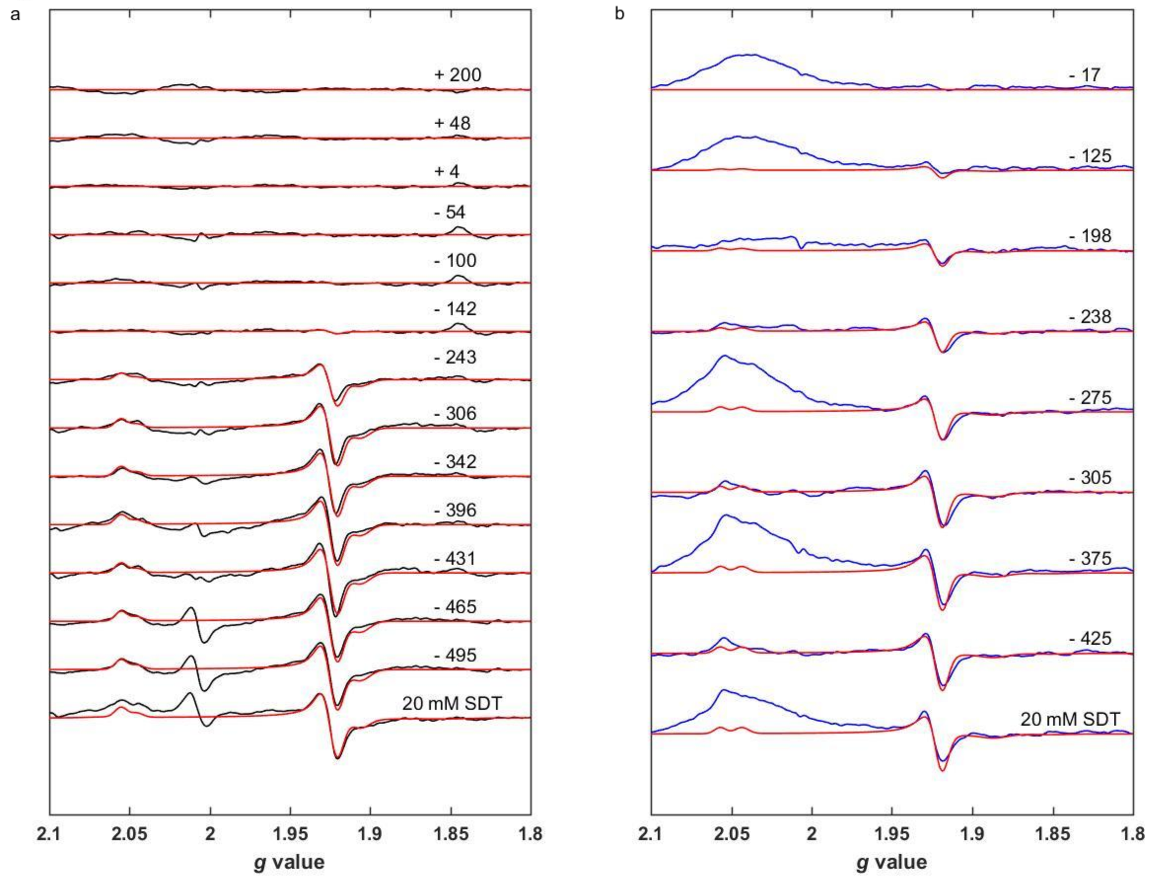

**Supplementary Fig 6**. Lack of N0 signal in *T. elongatus* PS-CI at −431 mV. EDFS spectra of *T. elongatus* titration sample at −431 mV measured at 10 K, 0.5 µs SRT, $\tau$ = 350 ns, $\pi$ = 32 ns. N2 simulated in pink, N1 simulated in purple (1:1), sum of simulation in red, with no indication that N0 is reduced.

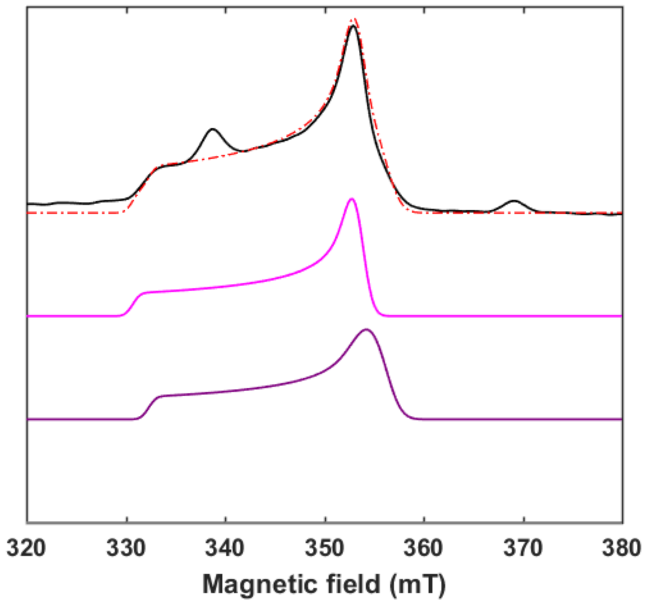
